# Harnessing Mass Spectrometry-Based Proteomics for Continuous Directed Evolution

**DOI:** 10.1101/2025.03.03.641153

**Authors:** Katharina Belt, David Obe, Mark A. Wilson, A. Harvey Millar, Ulschan Bathe

## Abstract

Continuous directed evolution is a powerful Synthetic Biology tool to engineer proteins with desired functions in vivo. Mimicking natural evolution, it involves repeated cycles of high-frequency mutagenesis, selection, and replication within platform cells, where the function of the target gene is tightly linked to the host cell’s fitness. However, cells might escape the selection pressure due to the inherent flexibility of their metabolism, which allows for adaptation. Whole-proteome analysis as well as targeted proteomics offer valuable insights into global and specific cellular changes. They can identify modifications in the target protein and its interactors to help understand its evolution and network integration. Using the continuous evolution of the Arabidopsis methionine synthases AtMS1 and AtMS2 as an example, we demonstrate how mass spectrometry-based proteomics can be applied in CDE, propose specific checkpoints for its integration and illustrate its role in informed decision making.

## INTRODUCTION

Directed evolution is one of Synthetic Biology’s most outstanding achievements. It aims at engineering biomolecules, e.g. enzymes, with required phenotypes or functions (1). Classical directed evolution was awarded the 2018 Nobel Prize (2), but continuous directed evolution (CDE) is considered even more powerful overcoming limitations in scale of experimentation and depth of evolutionary search. Thus, it holds the potential to shape future medicine, agriculture and industry.

The principle of CDE is *in vivo* sequence diversification, selection and replication, and couples the function of the target gene to the host cell’s fitness (3). Diverse CDE tools have been published that allow directed evolution in diverse platform cell types, including phage-based, bacterial, yeast or mammalian systems (4–10). With that, a wide range of targets can be engineered – from antibodies to substrate specificity of industrially relevant enzymes (11, 12); from improved catalytic efficiency to novel enzyme activities (13, 14); and from bacterial targets to genes from highly complex plants or mammals (15–17).

CDE is considered especially powerful because repeated cycles of mutagenesis and screening inside the platform organism allow efficient and scalable evolution. However, the host cells’ metabolism is a complex and interconnected protein machinery and CDE may cause trade-off effects in the proteome that can be easily overlooked during the evolution campaign. This might be caused by a selection pressure that is not tight enough or potentially affects other pathways; in the worst case, the platform cells escape the selection pressure entirely. In such cases, post-CDE evaluation of the engineered molecule would reveal that evolved phenotypes are not only or not exclusively connected to the target gene’s function. To reduce the risk of failure and to make CDE even more powerful, whole-system analysis might be used to look beyond single target genes, transcripts and metabolites. While whole genome and transcriptome sequencing would give some upstream insights into a cell’s real condition, metabolomics and proteomics deliver more information about the platform’s cellular changes because they are closer to the phenotype. They can directly show interaction of the engineered target with other biomolecules and pathways, and potential trade-offs. Proteomics is not utilized as a standard tool in SynBio experiments yet; for example, we are aware of only one case of a CDE study utilizing whole-cell proteomics (18). Mass spectrometry-based proteomics can provide a detailed snapshot of an organism’s entire proteome or of specific protein targets and their relative or absolute abundance, offering both qualitative and quantitative insights (19–21). It is also the method of choice to determine protein half-life and structural modifications due to its sensitivity and specificity. These attributes make proteomics an invaluable tool for CDE, enabling the verification of intermediate steps, validation of final functions, and troubleshooting of challenges during the evolution process. Here, we illustrate how proteomics can enhance CDE by identifying trade-offs and cellular adaptation during the continuous evolution of *Arabidopsis* methionine synthases AtMS1 and AtMS2.

## RESULTS AND DISCUSSION

### Host selection and validation for directed evolution

The selection of a platform host depends on the origin of the target gene and the destination organism, considering factors such as subcellular localization of the target protein, posttranslational modifications (PTM), intracellular pH, redox conditions, and oxygen levels. PCR and phenotypic validation (e.g., growth assays in defined media) are common techniques to validate the chosen background strain (e.g., a knockout mutant in *Saccharomyces cerevisiae*). However, complementing these analyses with targeted proteomics is advisable to confirm strain suitability and verify the presence of the target protein and absence of knocked out native protein (Figure 1, A). In addition, untargeted proteomics can reveal cellular changes that may influence selection during the evolution experiment.

**Figure 1.**
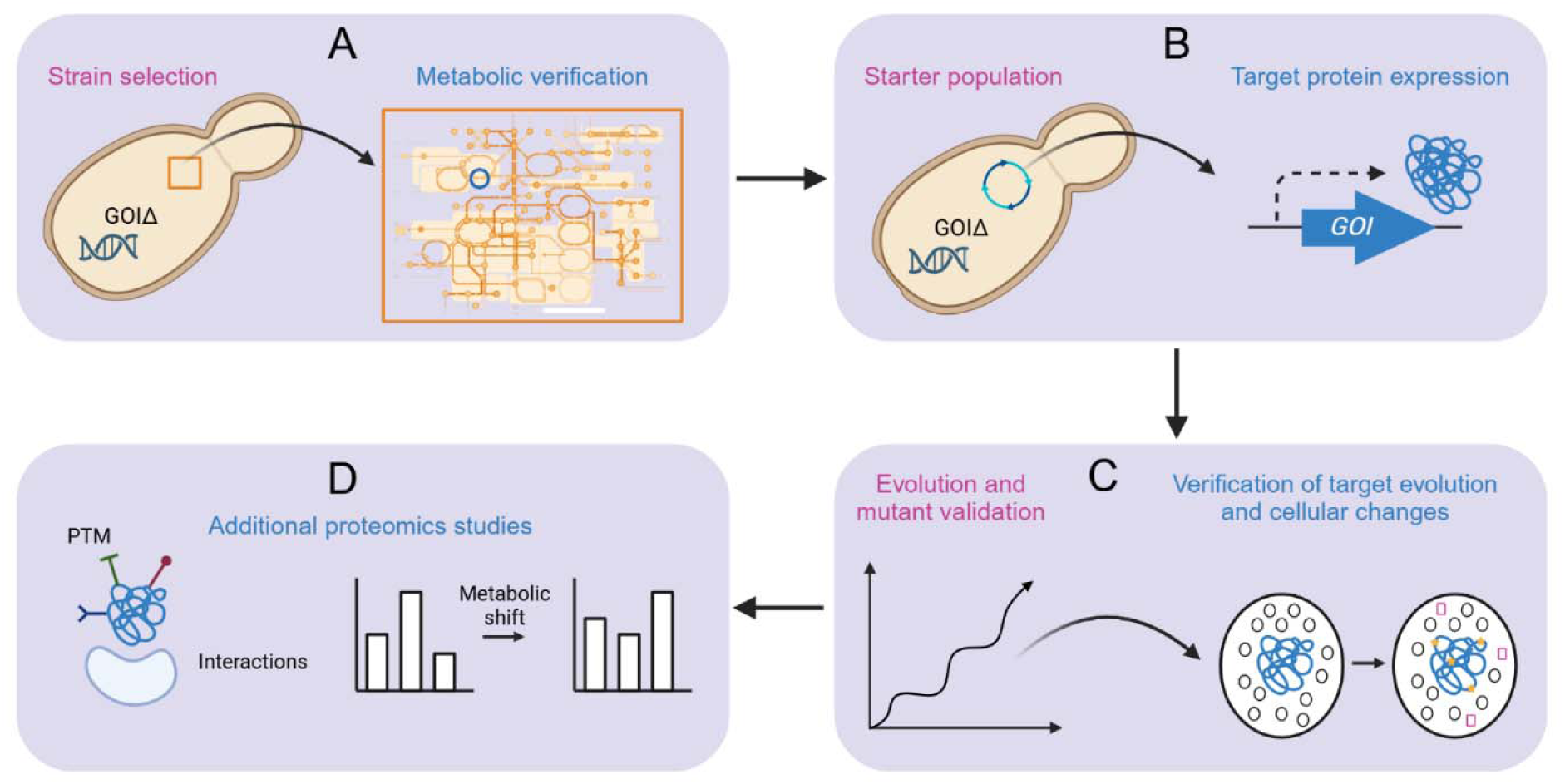
Proteomics in continuous directed evolution. This figure illustrates the experimental process of CDE with proteomic analysis incorporated at critical stages to evaluate protein-level changes (A-D). A, The CDE workflow begins with baseline proteomic analysis of the initial strain to assess global protein expression. B, Targeted proteomics can validate expression and abundance of the protein of interest. C, Random mutagenesis is applied to the gene of interest, followed by selection, with untargeted proteomic analysis identifying early protein shifts. Evolving strains are screened for phenotypic changes (e.g., growth), and proteomics verifies protein-level adaptations during evolution. The best-performing strains are finally selected, and proteomic validation confirms the protein function associated with the observed traits (D).

In this study, we aimed to evolve the short-lived methionine synthases AtMS1 and AtMS2 from *Arabidopsis thaliana* for longer working life (22, 23). We selected *Saccharomyces cerevisiae* as the evolution host due to its ‘plant-like’ cellular conditions, including compartmentalization, metabolite and cofactor availability, and shared metabolic pathways. For our approach, the AtMS1 and AtMS2 genes were codon-optimized for yeast and expressed from plasmid ArEc-TDH3 in a *met6*Δ knockout mutant to perform complementation assays in a methionine-deficient (-Met) medium (Supplemental Figure S1). Targeted proteomics can support this validation step if the desired phenotype is not achieved, enabling the assessment of i) protein abundance, ii) solubility, and iii) interactions with the cellular machinery.

### Generation of evolution starter populations and verification

When selecting a CDE system, factors such as mutation rate, the type of mutations introduced, and overall system design should align with the experiment’s objectives. For this study, we utilized the OrthoRep system due to its ability to achieve an orthogonal replication mutation rate between 10^5^ and 10^6^ times higher than the genomic mutation rate (7, 24). OrthoRep comprises an orthogonal error-prone polymerase expressed from a nuclear plasmid, and two cytosol-localized linear plasmids (p1 and p2) that facilitate self-replication, encode an RNA-polymerase, a marker gene and the gene of interest (Figure S2, A). To assemble this system, the gene of interest (here, AtMS1 or AtMS2) and the marker gene were first integrated into the p1 plasmid in the strain GA-Y319 (25). Successful integration was confirmed by estimating the p1 plasmid size on an agarose gel and by verifying the target gene via Sanger sequencing. In addition, we recommend that targeted proteomics analysis, as described earlier, is employed here to confirm the target protein’s expression and functionality in the OrthoRep setup.

To transfer the OrthoRep machinery to the chosen platform host, we performed protoplast fusion between the obtained donor strain carrying the p1 plasmid and the *met6*Δ recipient strain. Protoplast fusion clones were verified by gel analysis, Sanger sequencing of the target gene on p1 and complementation assays in –Met medium (Figure 2, B). Proteomics can be employed here to assess whether the yeast protein machinery, including the CDE system and the protein of interest, shows any unexpected alterations (Figure 1, B). For our study, we first performed targeted proteomics using multiple reaction monitoring mass spectrometry (MRM-MS) in starter populations to confirm the presence and accurately measure the abundance of AtMS1 and AtMS2 (Supplemental Figure S2). For that, we used spike-in standards of AtMS1 and AtMS2 that were expressed *in E. coli* and purified for MRM-MS experiments. Specific peptides (Supplemental Table 1) were selected prior to MRM-MS and used to identify AtMS1 and AtMS2 in total protein extracts. Using the abundance of spike-in standards we could perform absolute quantification to compare them and found that the AtMS1 and AtMS2 proteins were roughly equally expressed (between 1 and 2.5 fmol/µg total protein). We then used an untargeted proteomics approach to investigate broader host strain changes. Interestingly, the populations carrying AtMS1 showed downregulation of Met upstream biosynthetic and related enzymes (MET10, MET16, MET2, MET5, MET8, MET3) compared to AtMS2 cells (Figure 3). This illustrates that expression of foreign enzymes in a CDE system can have different impacts on the host cells’ metabolism and its adaptation, even though such enzymes share high sequence identity, e.g., AtMS1 and AtMS2 proteins are 92.81 % identical.

**Figure 2.**
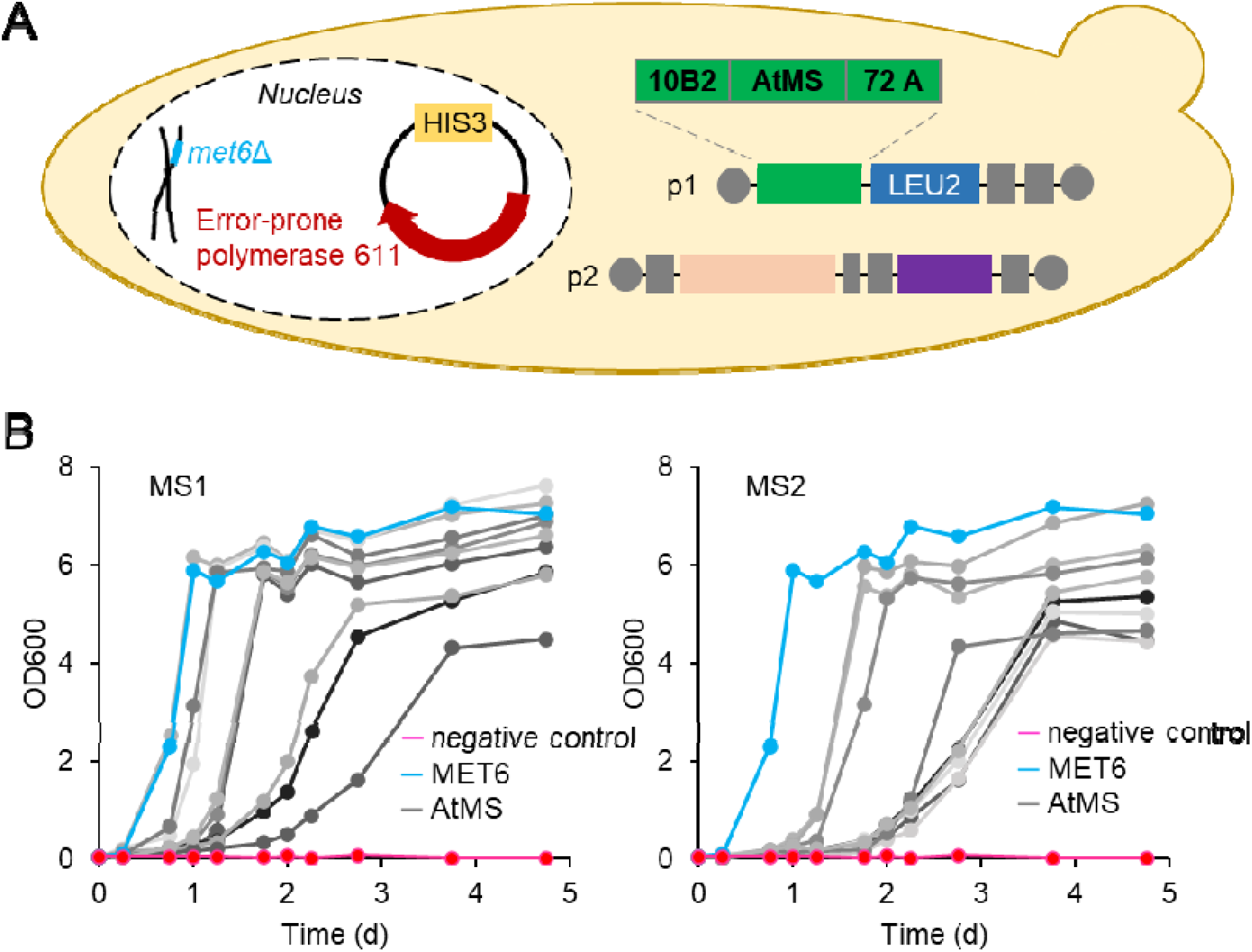
Assembly and test of evolution starter populations. A, Evolution populations were created in a BY4742 *met6*Δ yeast background. A nucleus-localized plasmid under control of a HIS3 marker encoded the error-prone polymerase TP-DNAP1_611. This polymerase is specific to genes on the linear, cytosolic plasmids p1 and p2. p1 encodes the MS target gene and a LEU2 selection marker. p1 and p2 harbor additional genes to sufficiently propagate the OrthoRep machinery. B, Complementation of BY4742 *met6*Δ with AtMS1 and AtMS2 expressed from p1 after protoplast fusion. Strains having the p1 with no MS gene served as negative control (magenta) but were supplemented with Met in the growth medium as positive control (blue). The MS populations were used to start the evolution campaign.

**Figure 3.**
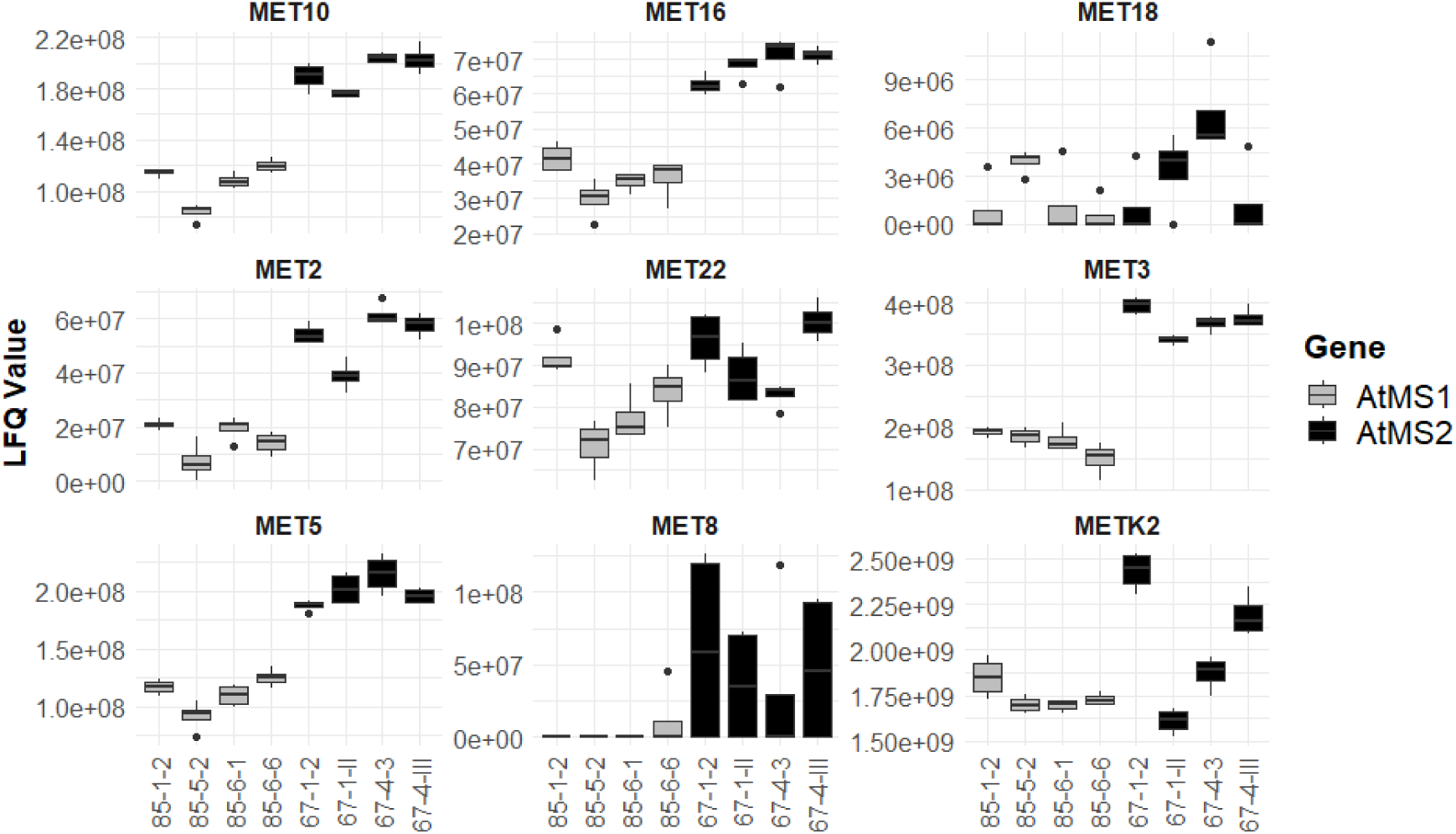
Protein analysis of Arabidopsis AtMS1 and AtMS2 in evolved and unevolved yeast populations. Untargeted proteomics analysis of protein abundance in the background of yeast host cells expressing either AtMS1 or AtMS2 from Arabidopsis. The figure shows Met synthase-related proteins in unevolved populations, with corresponding LFQ values obtained from mass spectrometry. Data include four replicates per strain.

### Monitoring ongoing evolution

Running CDE experiments are considered hands-off and low-effort ventures. Mutations in the target gene being beneficial are selected when the target’s performance is coupled to growth or another readable phenotype. Over time, these mutations accumulate, and the evolution is stopped when a desired phenotype is achieved. In practice, this means serial passaging, typically every couple of days, with growth phases in between. To evolve AtMS1 and AtMS2 for longer enzyme life, we used a selection strategy in which the concentration of the toxic Met analog L-selenomethionine (SeMet) in the medium was gradually raised from 5 µM to 1 mM (Supplemental Figure S3). This classical strategy favors increased Met synthase-driven flux to Met, which expands the free Met pool and so competes out the analog’s toxicity. Nine independent starting populations expressing either AtMS1 or AtMS2 were grown for 29–35 serial passages on Met-free medium containing SeMet. As growth improved, populations were typically split into two or more subpopulations to guard against accidental loss and to enable different evolutionary trajectories from that point forward (1). During the campaigns, we collected sequencing data of the target gene at passage 19 to verify that mutations accumulated. Such information may also help to follow the evolutionary trajectory. We observed mutations that replaced the wildtype base in all cells of the population as well as mixed sequences, i.e., not all copies of the target gene carried the mutation (data not shown). While such gene analysis is limited to sequence information, proteomics is a complementary tool to monitor the actual phenotype (Figure 1, C). For example, targeted proteomics can be used to assess changes in the abundance of the target protein and ensure its solubility and proper folding under the applied selection pressure. Additionally, untargeted proteomics can help identify global cellular responses to the selection regime, revealing potential off-target effects (e.g., genomic mutations) or adaptive mechanisms in the host (e.g., epigenetic adaptation). Such data can provide critical feedback to guide adjustments in experimental conditions and ensure that the desired phenotypic traits are being optimized effectively.

### Validation of evolved phenotypes

Once a desired phenotype is achieved, the evolution is stopped and evolved sequences are verified. We halted the MS campaigns at a final SeMet concentration of 1 mM as complex secondary toxic effects are more likely at higher concentrations (26, 27). In total, 72 populations were obtained. Figure 4, A shows the growth of six representative evolved populations compared to the corresponding unevolved populations in the presence of 1 mM SeMet. To verify increased intracellular Met concentrations as a result of longer MS life, we screened the 72 evolved populations in single replicates (Figure 4, B). Cellular free Met levels of 1–2 mM in most unevolved populations and a marked upshift in evolved populations towards levels as high as 10 mM pointed to successful evolution. Therefore, we chose eight and six populations with AtMS1 and AtMS2, respectively, to measure free Met pools in triplicates to gain statistical confidence about our data (Figure 4, C). Consistent with our previous screening, we confirmed high Met-producing and low Met-producing populations. Free Met pools may differ slightly from the first evaluation of populations due to the timepoint of harvest.

**Figure 4.**
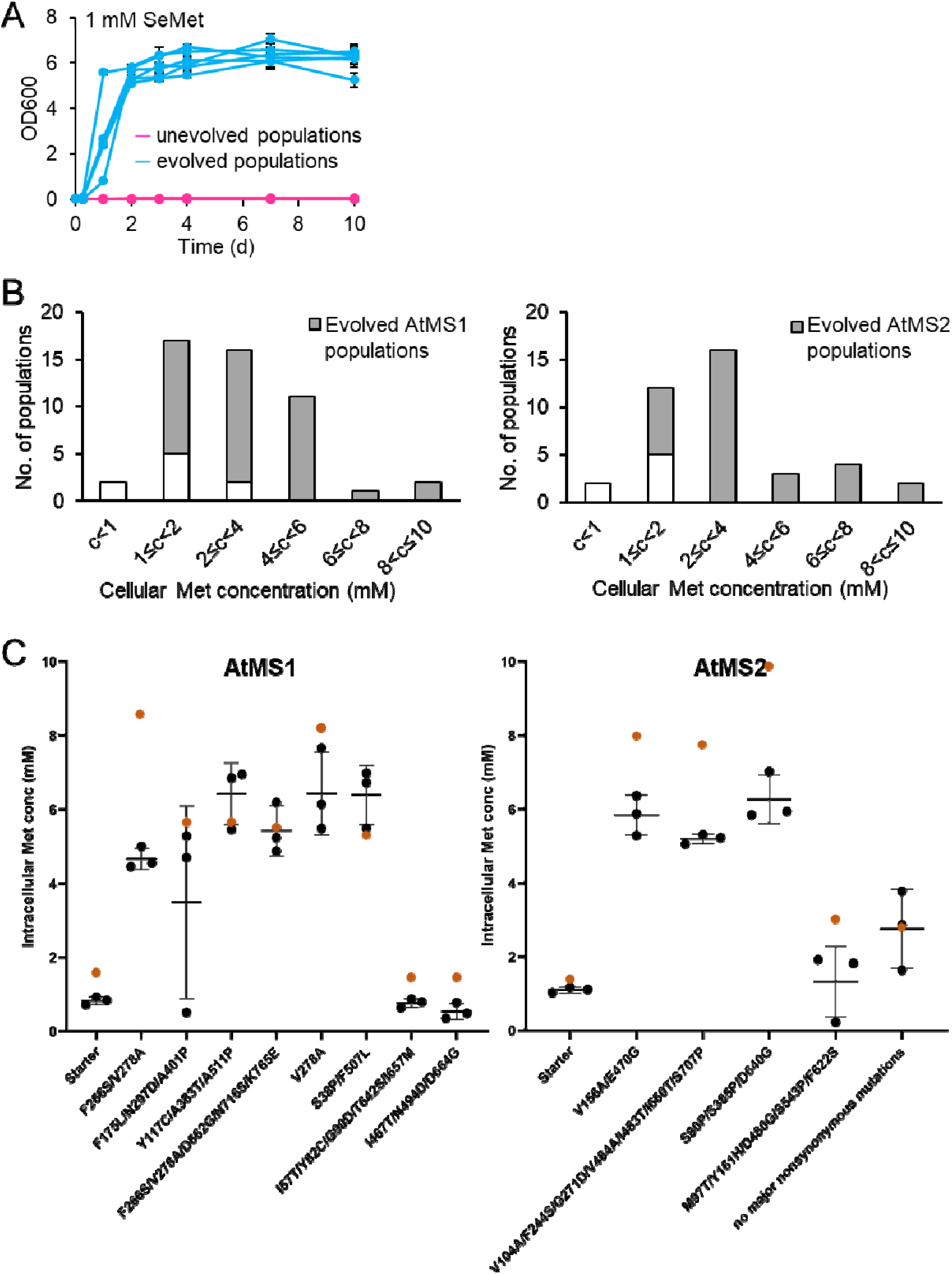
Analysis of evolved MS populations. A, Growth comparison of evolved (blue) and unevolved (cyan) populations in presence of 1 mM SeMet. B, Free intracellular Met pools of all unevolved (white) and evolved (grey) populations; n = 1. C, Evolved populations with the highest Met content from B were chosen for proteomics analysis and subjected again to free Met pool measurements; n = 3. Representative starter populations are given for comparison (left in each graph). Free Met pools from the first evaluation (B) of populations are given as orange dots.

To determine what mutations accumulated in CDE, the engineered sequence was isolated from the population and subjected to sequencing. The chosen MS populations from the Met pool measurements were used to detect mutations (Figure 5). Mutant MS variants were recovered having a mix of nonsynonymous, synonymous and promoter mutations. Most of them were considered ‘major’ mutations, i.e., they replaced the wildtype base in all gene copies of the population. They were truly selected and can be considered beneficial.

**Figure 5.**
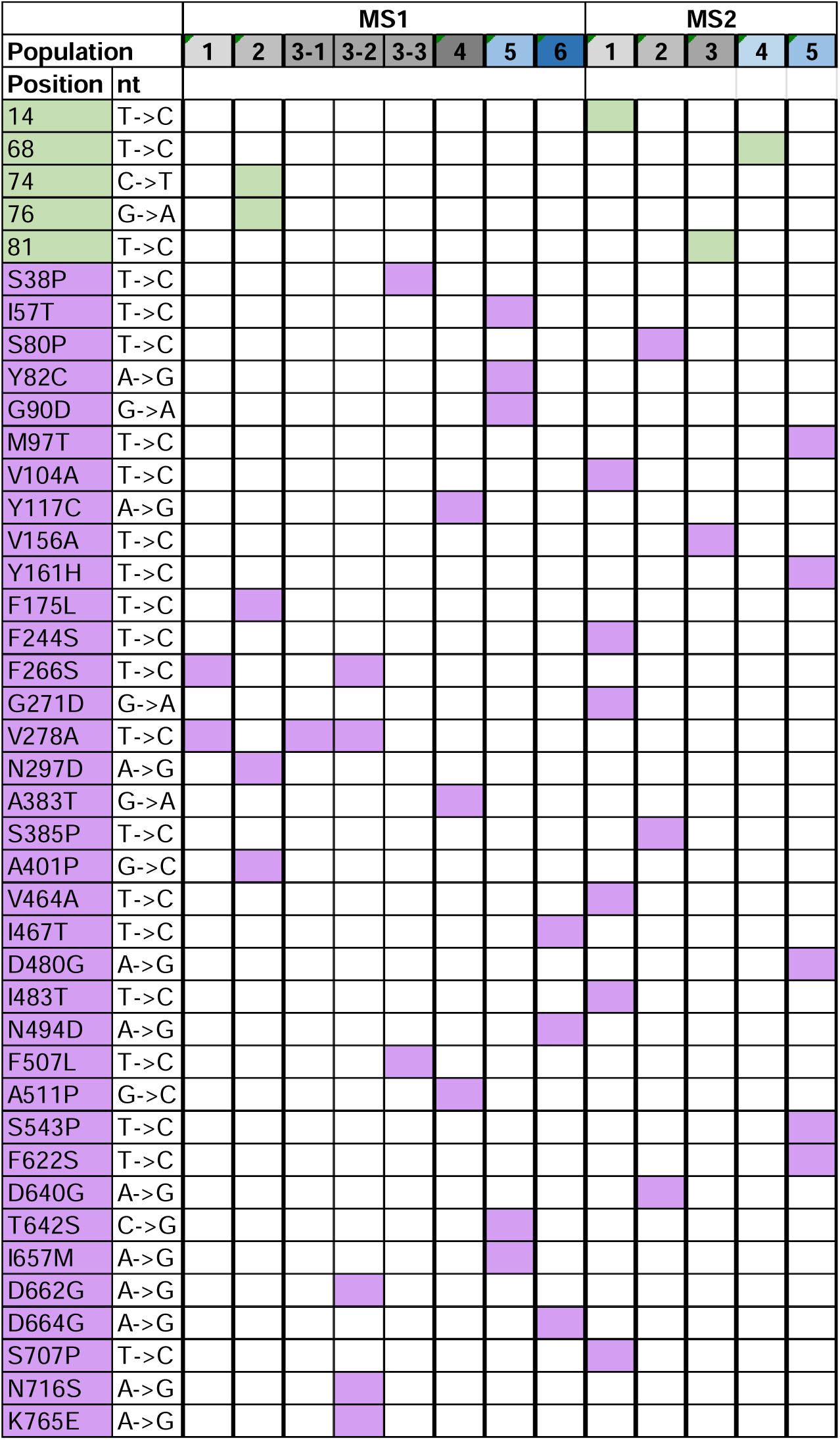
Major mutations recovered in evolved MS populations. Populations with high free Met pools chosen for further analysis (grey) accumulated promoter (green) and major ORF nonsynonymous (purple) mutations. Representative populations with low free Met pools (blue) are given for comparison.

As outlined above, proteomics can be applied to validate evolved phenotypes by directly assessing changes in abundance of proteins as outcomes of the evolution process in a targeted and untargeted manner (Figure 1, C). With that, broader cellular adaptations can be identified that might accompany the evolved phenotype, such as compensatory mechanisms or off-target effects. This comprehensive validation ensures that the evolved traits align with experimental objectives and that the evolved host functions as intended. We hypothesized that longer MS protein lifetime (i.e., highly protein stability) would correlate with higher protein abundance in evolved populations. Unexpectedly, using targeted proteomics, we measured lower MS abundances in evolved populations compared to unevolved ones, indicating that accumulated major mutations did not correlate with evolved cellular phenotypes, e.g. elevated free Met pools and improved growth in medium with SeMet (Figure 6, A). Nonetheless, the proteomics approach demonstrated sensitivity and statistical confidence, enabling the detection and quantification of even very low protein abundances, down to femtomoles per microgram of the yeast background proteome.

**Figure 6.**
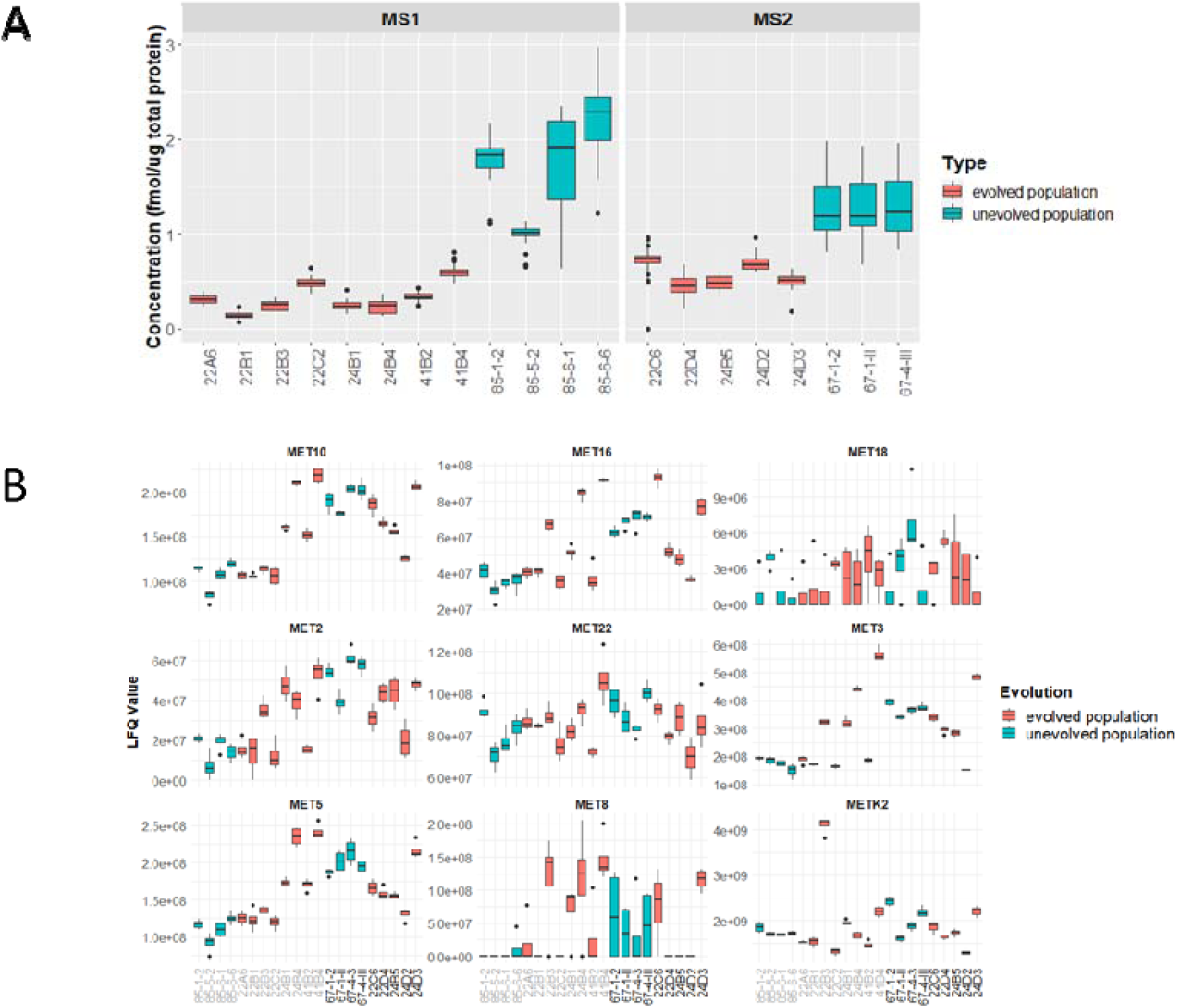
Protein analysis of Arabidopsis AtMS1 and AtMS2 in evolved and unevolved yeast populations. A, Concentrations of Arabidopsis AtMS1 and AtMS2 proteins in evolved and unevolved yeast populations, quantified by mass spectrometry. Data represent the median protein concentrations from four replicates per yeast strain. B, Untargeted proteomics analysis of both evolved and unevolved populations expressing either AtMS1 (grey labels) or AtMS2 (black labels) from Arabidopsis. Expression of Met synthase-related proteins with corresponding LFQ values obtained from mass spectrometry are shown. Data include four replicates per strain.

We then used untargeted proteomics to investigate broader host strain changes, in particular in the Met biosynthesis and related pathways. This analysis identified alterations in the evolved populations for which we could not find correlations with either free Met pools, the target gene (AtMS1 or AtMS2) nor accumulated mutations (Figure 6, B; Supplemental Table 2). It demonstrates that the applied selection pressure influenced not only the target protein (i.e., major mutations accumulated) but also affected the host’s metabolic network. These findings highlight the complementary strengths of targeted and untargeted proteomics for a comprehensive understanding of evolved phenotypes and system-wide effects.

### Beyond Phenotype Validation: Additional Proteomics Approaches

To further enhance the insights gained from CDE, proteomics analyses could also be integrated at others stages in the process (Figure 1, D). First, post-translational modification analysis can provide valuable information on how target proteins are regulated or modified in the host or during CDE, potentially revealing novel mechanisms of adaptation. For example, phosphorylation, acetylation, oxidation or glycosylation patterns being added or deleted during evolution could impact protein activity and interactions in the host environment. Second, interaction proteomics, such as co-immunoprecipitation coupled with mass spectrometry, can be employed to identify changes in protein-protein interaction networks that involve the evolved proteins. This is particularly useful for understanding how the evolved phenotype integrates into the cellular machinery. Thirdly, proteomics-based quantification of metabolic enzymes can link phenotypic changes to metabolic shifts, providing a more holistic understanding of how evolution reshapes cellular physiology. These additional analyses can deepen the mechanistic understanding of evolved phenotypes and inform strategies for optimizing the evolutionary process further.

## CONCLUSION

CDE experiments greatly benefit from implementing mass spectrometry-based proteomics because it can uncover cellular changes which complement what could be detected on the DNA or RNA levels. As shown in our study, it can give valuable insights into the actual phenotype of the platform cells during a campaign. Using *Arabidopsis* AtMS1 and AtMS2 as an example, we demonstrated at what checkpoints both targeted and untargeted proteomics can be applied, what the delivered value would be, and outlined other potential applications during and after evolution. We discovered that different target proteins that were subjected to evolution can have varying effects on the host cell, even though they share high sequence similarity. Furthermore, we identified changes in protein expression of upstream and related metabolic pathways as a result of the applied selection. Combined, our results demonstrate the benefits of complementing continuous evolution with proteomics analysis.

## ASSOCIATED CONTENT

### Supporting Information

#### MATERIAL AND METHODS

Supplemental Figure S1. Complementation of BY4742 met6Δ with AtMS1 and AtMS2.

Supplemental Figure S2. Targeted protein analysis of Arabidopsis AtMS1 and AtMS2 in evolutions starter populations.

Supplemental Figure S3. SeMet treatment to evolve MS.

Supplemental Table S1. Peptide sequences and mass spectrometry analysis parameters used for targeted selected reaction monitoring analysis of AtMS1 and AtMS2.

Supplemental Table S2. Untargeted proteomics of evolved and unevolved yeast strains with either AtMS1 or AtMS2 from *Arabidopsis*.

Supplemental Table S3. Sequences of genes used in this study.

Supplemental Table S4. Primers used for amplification and sequencing.

## AUTHOR INFORMATION

### Corresponding Author

*Ulschan Bathe

### Author Contributions

U.B., K.B. and A.H.M. conceived and designed the project; U.B. performed directed evolution and metabolite quantification; D.O. and M.A.W. expressed and purified proteins as standards for proteomics; K.B. and A.H.M. performed proteomics analyses; K.B. and U.B. wrote the article with contributions from other authors. U.B. agrees to serve as the author responsible for contact and ensures communication.

## ACKNOWLEDGMENT

We thank Andrew D. Hanson from the University of Florida for his support as well as his valuable advice and guidance. We thank Elke Ströher from the WA Proteomics Facility at the University of Western Australia for her support with mass spectrometry analysis.

## ABBREVIATIONS

CDE: continuous directed evolution
PTM: posttranslational modifications
AtMS1: *Arabidopsis thaliana* methionine synthase 1
AtMS2: *Arabidopsis thaliana* methionine synthase 2
MRM-MS: multiple reaction monitoring mass spectrometry

## BRIEFS

Mass spectrometry-based proteomics was leveraged to enhance an OrthoRep directed evolution strategy.

