## Supplemental Information for "Harnessing Mass Spectrometry-Based Proteomics for Continuous Directed Evolution"

**for**

**MATERIAL AND METHODS**

**Plasmid construction**

Gene sequences are given in Supplemental Table S3. Primer sequences used for amplification and sequencing were purchased from Eurofins Genomics (Louisville, KY) and are listed in Supplemental Table S4. Non-recoded cDNAs of *Arabidopsis* AtMS1 (AT5G17920) and AtMS2 (AT3G03780) were obtained from the ABRC stock center (<https://abrc.osu.edu/>), G13034 and C104738. *MET6* was amplified from genomic DNA of *Saccharomyces cerevisiae* strain BY4742. ORF of AtMS1, AtMS2 and MET6 were cloned into plasmids ArEc-TDH3 and GR-306MP as previously described (1). Enzymes for PCR and cloning were obtained from Thermo Fisher Scientific (Miami, FL).

**Yeast strain construction and media**

Yeast strain BY4742 *met6*Δ (MATα; his3Δ1; leu2Δ0; lys2Δ0; ura3Δ0; YER091c::kanMX4) was obtained from Euroscarf (Oberursel, Germany), and grown in YPD medium as previously described (1). GA-Y319 was cultivated as described previously (1). Plasmid ArEc-TDH3 containing AtMS1, AtMS2 or MET6 were transformed into BY4742 *met6*Δ using the lithium acetate (LiAc) method with minor modifications (2). A 5-mL YPD preculture was inoculated with BY4742 *met6Δ* and grown to saturation at 30°C. Five milliliters of YPD per transformation was inoculated with the preculture to set an OD_600_ of 0.3 and grown to an OD_600_ of 0.6. Cells were pelleted by centrifugation (5 min, 1,500 g), washed with 5 mL of Milli-Q water per 5 mL of culture, recentrifuged, washed with 150 µL of TE/LiAc (10 mM Tris–HCl, 1 mM Na_2_EDTA, pH 7.5, 100 mM LiAc) and then mixed with 50 µL of TE/LiAc per 5 mL of culture. Ten milliliters of 11 mg/mL of single-stranded salmon sperm DNA were mixed with 300 ng of plasmid DNA, and added to the cells. After adding 600 µL of 50% w/v PEG 3350 plus 100 µM of LiAc and 100 µM of TE (10 mM Tris–HCl, 1 mM Na_2_EDTA, pH 7.5), the mix was incubated for 45 min at 30°C with agitation, and afterwards for 20 min at 42°C. Cells were then pelleted by centrifugation (5 min, 700 g) and resuspended in 1 mL of Milli-Q water. Selection was done in synthetic complete (SC) minus histidine for 3 d at 30°C. Selection media were SC-His (6.7 g/L YNB with ammonium sulfate, 1.4 g/L Dropout mix synthetic minus histidine, 20 g/L Bacto Agar (when needed), 2% w/v glucose) for cells containing ArEc-TDH3 plasmids, SC-Leu (6.7 g/L YNB with ammonium sulfate, 1.4 g/L Dropout mix synthetic minus leucine, 20 g/L Bacto Agar (when needed), 2% w/v glucose) for GA-Y319 containing p1_MS, and SC-His-Trp-Leu-Met-Cys (6.7 g/L YNB with ammonium sulfate, 1.4 g/L Dropout mix synthetic minus histidine, tryptophan, leucine, methionine and cysteine, 20 g/L Bacto Agar (when needed), 2% w/v glucose) for BY4742 *met6*Δ containing p1_MS. Strains designed for validation of mutant variants were grown in SC (6.7 g/L YNB with ammonium sulfate, 1.4 g/L Dropout mix synthetic minus histidine, tryptophan, uracil, methionine and cysteine, 20 g/L Bacto Agar (when needed), 2% w/v glucose). Transformation of GA-Y319 was done with ScaI-digested GR-306MP harboring AtMS1 or AtMS2 as previously described (3). Single colonies were selected for total DNA isolation and sequencing to confirm integration of the MS genes and the leucine selection marker in the p1 plasmid (1). Primers used are listed in Supplemental Table S4. Sequence-verified clones were used as the donor strain for protoplast fusion and BY4742 *met6*Δ as recipient. Protoplast fusions were made as before (1). Single colonies were selected for total DNA isolation and sequencing to confirm integration of the MS mutant variants and the uracil selection marker in the p1 plasmid (1). Primers used are listed in Supplemental Table S4.

**Complementation and growth assays**

For complementation assays, single colonies of BY4742 *met6*Δ transformed with ArEc-TDH3 containing AtMS1, AtMS2 or MET6 were grown in 5 mL SC-His at 30°C until saturation. Cells with ArEc-TDH3 only were used as negative control. An aliquot of the cultures was used to inoculate triplicates of 5 mL SC-His at a starting OD_600_ of 0.05. Cultures were grown at 30°C for 5 d and the OD_600_ was monitored. Growth assays were done similarly but in SC medium minus methionine, cysteine, histidine, tryptophane and leucine plus 1 mM SeMet.

**Evolution campaigns**

Three-milliliter cultures were started at OD_600_ = 0.05 in SC minus methionine, cysteine, histidine, tryptophane and leucine, and cultivated in Corning deep 24-well plates (Millipore Sigma, catalog no. AXYPDW10ML24C). Plates were covered with Breathe-Easier Sealing Film, Diversified Biotech (Dedham, Massachusetts). Cultivation was performed at 30°C with humidification at 80% and agitation at 800 rpm in an Infors Multitron shaker (Sulzemoos, Germany). SeMet was added from the start at a concentration of 5 µM. Populations were subcultured into fresh medium when they reached saturation. SeMet concentration in the medium was ramped up whenever the evolving populations reached saturation after two days of cultivation. Evolving populations were split at passages 6, 13, and 29. Bulk cultures were analyzed for mutations by sequencing PCR amplicons containing the MS ORF and promoter.

**DNA isolation and sequencing**

Five-milliliter cultures of evolved populations were used for total DNA extraction as previously described or as follows (1). Cells from a 100 µL aliquot of cultures were mixed with 100 µL of 200 mM lithium acetate plus 1% SDS, and incubated for 10 min at 70°C. 300 µL of ethanol were added, mixed and centrifuged for 3 min at 15,000 x g. The pellet was washed with 500 µL of 70% ethanol and centrifuged again. After drying for 5 min at 50°C, the pellet was resuspended in 100 µL of milliQ water and centrifuged again for 15 sec. 90 µL were then transferred to a new tube and used as template for PCR amplification and subsequent Sanger sequencing. Primers used are listed in Supplemental Table S4.

**Analysis of free amino acid pools**

Precultures of unevolved and evolved populations were grown in 3 mL SC minus histidine, tryptophane and leucine, and cultivated until saturation in Corning deep 24-well plates as described above. Aliquots of precultures were inoculated into 3 mL SC minus methionine, cysteine, histidine, tryptophane and leucine at an OD_600_ of 0.05, and cultivated as before until they reached late log phase. Per culture, a liquid amount, that equals around 300 Mio. cells, was harvested by centrifugation, and the cells were stored at -80°C until extraction. Free amino acids were extracted from the yeast cultures as described previously with the following modifications (4). 500 µL of 12:5:1 methanol-chloroform-water were added to the frozen cells and mixed thoroughly. Three separate samples were used to determine the spike recovery. They were split into two each and one aliquot per sample was mixed with 50 µL of a 2.5 mM methionine solution. The downstream procedure was as for all other sample but with half of the used volumes. The samples were incubated for 5 min at 50°C, mixed thoroughly again and centrifuged for 1 min at 21,000 x g. The supernatant was transferred to a new tube and the remaining cells were extracted again. 250 µL of chloroform and 375 µL of Milli-Q water were added to the pooled supernatants and mixed by inverting the tubes. The mix was centrifuged for 5 min at 21,000 x g, and 1 mL of the aqueous upper phase was recovered and dried in a speed vac. Derivatization of amino acids was done as described previously with minor changes (4). The dried extracts were resolved in 100 µL of Coupling solution (10:5:2:3 Acetonitrile: Pyridine: Triethylamine: H_2_O), dried again, resolved in 100 µL of Coupling solution and mixed with 5 µl PITC (Edman's Reagent; phenylisothiocyanate) (Thermo Fisher Scientific). The solution was then incubated for 5 min at room temperature, dried, and the extract was finally resolved in 125 µL Sample buffer (0.071 g anhydrous sodium dihydrogen phosphate in 100 mL of Milli-Q water, adjusted pH to 7.4, mixed with 5.26 mL acetonitrile). Samples were centrifuged before HPLC analysis. Reverse-phase HPLC analysis of the free methionine pool was carried out as described (5) using a Waters^TM^ (Milford, MA) 2695 Separations Module coupled to a Waters^TM^ 2998 PDA Detector. Chromatographic separation was performed on a Supelco^TM^ Discovery® C18 column (Merck KGaA, Darmstadt, Germany) at 25°C. The injection volume was 5 µL. The flow rate was 1 mL/min, and the gradient was as follows. 10 min at 100% eluent A (131 mM sodium acetate, 3.38 mM triethylamine, adjusted to pH 5.7 with acetic acid, v/v 0.06% acetonitrile, 2.4 µM EDTA), 30 s at 46% eluent B (3:2 acetonitrile:water), 2 min at 100% B, 8 min at 100% A. Detection was done at a wavelength of 254 nm. Met identification was done based on comparison with an authentic standard (Merck KGaA, Darmstadt, Germany), and concentrations were calculated based on a standard curve and spike recovery rates.

**Protein expression in *E. coli* and purification**

For spike-in standard, *Arabidopsis thaliana* AtMS1 and AtMS2 genes were synthesized with codon-optimization for expression in *E. coli* and then cloned into pET-15b vector at the NdeI and XhoI restriction sites. The pET-15b-AtMS1 and pET-15b-AtMS2 constructs were expressed in *E. coli BL21(DE3*) using standard protocols as previously described (6). Briefly, cells were cultured at 37 °C with shaking in LB medium containing 100 µg/mL ampicillin and supplemented with 0.5 mM ZnSO_4_ until the OD_600_ of the culture reached 0.6. Protein overexpression was induced by adding filter-sterilized isopropyl β-D-1-thiogalactopyranoside (IPTG) to a final concentration of 0.2 mM for AtMS1 and 0.4 mM for AtMS2. The culture was cooled and cells were grown with shaking at 18 °C overnight and then treated with 100 µg/ml of chloramphenicol one hour prior to harvesting to help recover soluble protein from inclusion bodies (7). Cells were collected via centrifugation at 4°C and stored at −80 °C.

Harvested cells were resuspended in lysis buffer (300 mM sodium chloride, 50 mM HEPES pH 7.4). The cells were then incubated with 1 mg/ml lysozyme at 4°C for 1 h and then lysis was completed via sonication. The crude lysate was centrifuged (20 min, 30,000 × *g*, 4 °C) and the pellet was discarded. The clarified supernatant was mixed with Ni^2+^-NTA resin (Millipore Sigma, cat. no. P6611) and incubated at 4°C on a nutator for 30 min. The slurry was used to pour a column and AtMS1/2 was eluted using elution buffer (300 mM sodium chloride, 50 mM HEPES pH 7.4, and 250 mM imidazole pH 7.0). The eluted MS proteins were incubated with thrombin (MP Biomedicals, cat. no. 154163) at a ratio of 1.5 U/mg of protein and then dialyzed against 1 L of 50 mM ammonium bicarbonate at 4°C overnight. The protein was passed through a Ni^2+^-NTA column to remove any protein that still retained the hexahistidine tag, followed by benzamidine-sepharose resin (Cytiva, cat. no. 17512310) to remove thrombin. Protein purity was determined using Coomassie-stained SDS-PAGE. The purified MS proteins were concentrated to 17 mg/ml as determined by UV-visible spectrophotometry using extinction coefficients at 280 nm (ε_280_) calculated by ProtParam (Expasy) (8) and then flash-frozen in 50-100 µL aliquots in liquid nitrogen for storage at -80°C. These were used to verify peptides for targeted MS analysis and as spike-in standards to allow absolute quantification.

**Cultivation and harvest of evolution populations for proteomics analysis**

Evolved yeast populations and their corresponding starter populations were grown in selective medium in Corning deep 24-well plates as described above. Per population, twelve cultures á 3 mL were harvested at an OD_600_ between 2.5 and 3.7 and pooled. The cells were harvested by centrifugation at 4°C and washed with 15 mL ice-cold milliQ water. The final pellet was resuspended in 1 mL ice-cold milliQ water, flash-frozen in liquid nitrogen and freeze-dried using a Labconco™ FreeZone™ 2.5L -50°C Benchtop Freeze Dryer (Catalog no. 710201000).

**Sample preparation for mass spectrometry analysis**

Proteins of interest, methionine synthases AtMS1 and AtMS2, were analyzed from purified proteins and dried yeast extracts, including 84 protein samples derived from unevolved and evolved populations.

Protein extracts were prepared by incubating 10 mg dried yeast samples in 200 µl SDS extraction buffer (7% SDS, 0.1 M Tris, 5 mM TCEP, 1x protease inhibitor, pH 7.5), followed by 10 min incubation at 95°C and 1000 rpm. Samples were transferred for a 1-minute treatment in a sonicator water bath to enhance lysis. The samples were centrifuged for 10 min at max speed (~18000 g), and the supernatant was transferred into a 96-well plate for further processing.

**Protein quantification and digestion**

Protein quantification was performed using a bicinchoninic acid (BCA) assay to estimate protein concentration. Proteins were digested using an SP3-aided protocol (9), employing trypsin at a 1:50 enzyme-to-substrate ratio. Digestion was carried out at 37°C for 18 h to ensure complete cleavage. From each sample, 10 µL of the prepared solution was injected for analysis.

**Mass spectrometry analysis**

For untargeted analysis, protein samples were injected into an online nanoflow system (0.4 µL/min) using a capillary column (Picofrit, 50 µm tip opening, 75 µm diameter, New Objective, ICT36015030F-50) packed in-house with 15 cm of C18 silica material (3 µm, Dr. Maisch GmbH). This was connected to a Thermo Fusion mass spectrometer in-line with a Dionex Ultimate 3000 series UHPLC. The spray voltage was set to 2 kV, and the heated capillary temperature was maintained at 275°C. For data-dependent acquisition (DDA) experiments, the full MS resolution was set to 60,000 at *m/z* 200, with an AGC target of 300% and auto injection time. The mass range was set from 350 to 1500 *m/z*. The AGC target value for fragment spectra was set to ‘standard’ with a resolution of 15,000 and auto injection time. An intensity threshold of 5 × 10^4^ was applied, and the isolation width was set at 1.6 *m/z*. Normalized collision energy was set to 30%.

Raw files were processed using the MaxQuant software (Version 2.4.0.0, (10)) using default settings. Trypsin was used as protease and up to two allowed missed cleavages. Cysteine carbamidomethylation was set up as a fixed modification and oxidation of methionine and N-terminal acetylation as a variable modification. Peptide identification required a minimum of seven peptides and a maximum of five modifications was allowed. Match-between-run function was enabled. Global data normalization was done using the MaxQuant LFQ algorithm with LFQ minimum ratio count set to 2 and fast LFQ enabled. The following fasta files were used: sequences for the evolved proteins plus the reference fasta file UP000002311 for Saccharomyces cerevisiae was downloaded from Uniprot. MaxQuant output files were used to further analyze and visualize in R.

For targeted analysis by multiple reaction monitoring, digested peptides were loaded onto an Aeris 3.6 µm peptide XB-C18 100 Å, LC column 50 x 2.1 mm (Phenomenex) using a Thermo UltiMate 3000 RSLCnano System coupled to an Thermo TSQ Altis Triple Quadrupole MS. The column was heated to 55°C and the column flow rate was 0.4 mL·min^-1^ with the following elution profile: 0–20.5 min 2% (v/v) acetonitrile, 0.1% (v/v) formic acid to 22% (v/v) acetonitrile, 0.1% (v/v) formic acid; 20.5–27.5 min 22% (v/v) acetonitrile, 0.1% (v/v]) formic acid to 35% (v/v) acetonitrile, 0.1% (v/v) formic acid; 27.5–28.5 min 35% (v/v) acetonitrile, 0.1% (v/v) formic acid to 97% (v/v) acetonitrile, 0.1% (v/v) formic acid; 28.5–29.2 min 97% (v/v) acetonitrile, 0.1% (v/v) formic acid, 29.2–29.5 min 97% (v/v) acetonitrile, 0.1% (v/v) formic acid to 2% (v/v) acetonitrile, 0.1% (v/v) formic acid; 29.5–35 min 2% (v/v) acetonitrile, 0.1% (v/v) formic acid. The list of peptide transitions used for selected reaction monitoring (SRM-MS) is provided in Supplementary Table S1. Peak area of ions for targeted peptides was determined using the Skyline software package version 24.1.0.199 (reference for skyline: https://pubmed.ncbi.nlm.nih.gov/20147306/). Further data analysis was performed using R.

**SUPPLEMENTAL FIGURES**

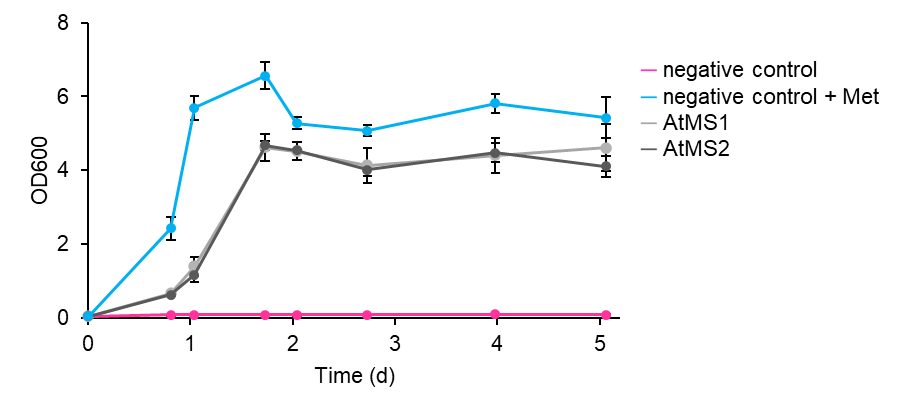

**Supplemental Figure S1: Complementation of BY4742 *met6*Δ with AtMS1 and AtMS2.** Proteins AtMS1 or AtMS2 were expressed from the plasmid ArEc-TDH3. The vector alone (i.e., no MS gene) served as negative control (magenta) but was supplemented with Met in the growth medium as positive control (blue).

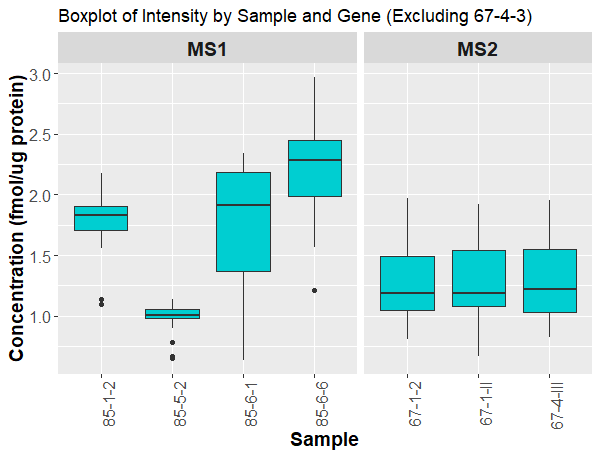

**Supplemental Figure S2: Targeted protein analysis of Arabidopsis AtMS1 and AtMS2 in evolutions starter populations.**

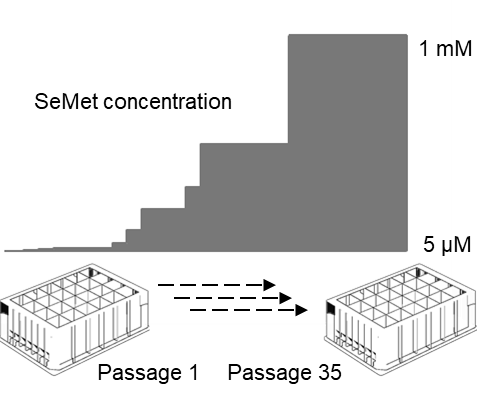

**Supplemental Figure S3: SeMet treatment to evolve AtMS.** AtMS evolution campaigns were run for 29–35 passages, and SeMet concentration was gradually raised from 5 µM to 1 mM.
